# Endogenous RNA editing of a nuclear gene *BOSS* triggers flowering in tomato

**DOI:** 10.1101/2021.06.09.447702

**Authors:** Wenqian Wang, Jie Ye, Chuying Yu, Qingmin Xie, Xin Wang, Huiyang Yu, Jianwen Song, Changxing Li, Long Cui, Heyou Han, Changxian Yang, Hanxia Li, Yongen Lu, Taotao Wang, Yuyang Zhang, Junhong Zhang, Bo Ouyang, Zhibiao Ye

## Abstract

RNA editing is defined as the production of transcripts with RNA sequences different from those of the DNA template. Most of previous RNA editing studies have focused on organelles, while RNA editing of nuclear transcripts is largely unknown. Here, we describe the first example of nuclear transcript RNA editing in plant, the *BOSS* RNA editing regulates tomato flowering. The SNP (SNP1) located in the 5’UTR of *BOSS* gene (the <u>B</u>alancer <u>o</u>f *<u>S</u>P* and *<u>S</u>FT*) is associated with tomato early flowering, and two transcripts of *BOSS* produced by SNP1 associated RNA editing show functional differentiation, where *BOSS*-β transcript promotes flowering while *BOSS*-α does not. Furthermore, these two transcripts of *BOSS* are shown to regulate *SP* (anti-florigen) pathway at transcription level in the shoot apical meristem (SAM). Our findings reveal a new layer of complexity in the control of plant stem cell proliferation and provide the evidence of RNA-editing of a single gene for flowering, suggesting that molecular breeding programs to increase the RNA-editing efficiency may improve the productivity of tomatoes and other agricultural organisms.

## INTRODUCTION

Crop yields are strongly associated with flowering time, and for many plants, the significance of flowering mainly involves two aspects: (i) balancing and optimizing their market supply for human demand, and (ii) enhancing their adaptability and expanding the geographical range of cultivation (Andres and Coupland, 2012; Wang et al., 2016). Tomato (*Solanum lycopersicum*), a main vegetable crop originating in the Andes of South America with a worldwide distribution, is classified as a day-neutral and temperature-loving plant, and its flowering and fruiting can be restricted under adverse conditions (Calvert, 1959; Song et al., 2013; Soyk et al., 2017).

Flowering is a sign for the transition of plants from vegetative to reproductive growth, and this process is precisely regulated by both genetic factors and external environmental factors (Imaizumi and Kay, 2006; Jung and Muller, 2009; He, 2012). In the long-day plant *Arabidopsis*, flowering has been shown to be controlled by five pathways, including vernalization, photoperiod, gibberellin, autonomous and senescence pathways (Srikanth and Schmid, 2011). These five pathways were integrated into a network through three central genes: *CO* (*CONSTANS*) (Suarez-Lopez et al., 2001; Valverde et al., 2004), *FT* (*FLOWERING LOCUS T*) (Kardailsky et al., 1999; Tiwari et al., 2010), and *FLC* (*FLOWERING LOCUS C*) (Michaels and Amasino, 1999; Searle et al., 2006). In the short-day plant rice, several genes have been identified to control the flowering time, such as *Hd3a (Heading date3a)* and its paralog *RFT1* (*RICE FLOWERING LOCUS T1*) encoding florigens (Kojima et al., 2002; Komiya et al., 2008; Komiya et al., 2009).

Despite increasing reports of single genes responsible for the variation of important traits like plant flowering (Teo et al., 2014), little is known about the synergistic regulation of flowering by different flowering genes or one gene through different transcriptional or post-transcriptional modification. In *Arabidopsis*, FLM-β and FLM-δ, two FLM protein splice variants with opposite functions, are shown to compete for interaction with the floral repressor SVP to dynamically regulate flowering at different temperatures (Pose et al., 2013). In sympodial plants such as tomato (*Solanum lycopersicum*) (Knapp et al., 2004; MacAlister et al., 2012), the differentiation of the first floral meristem occurs after the vegetative growth of several leaves, which determines the flowering time of tomato, termed as the first inflorescence node (FIN), followed by the vegetative growth from the axil of the youngest leaf through a sympodial vegetative meristem (SYM) to three leaves and then transition into a flower (MacAlister et al., 2012). Sympodial cycling is regulated by the balance between flower-promoting (florigen) and flower-repressing (anti-florigen) activities in the shoot apical meristem (SAM), with *SP* as a flowering repressor to transform tomato indeterminate shoot architecture into determinate vines (Pnueli et al., 1998), while *SFT* as a genetic originator of the flowering hormone florigen to promote flowering (Lifschitz et al., 2006). The dose-sensitivity balance of *SFT* and *SP* transcripts in SAM determines their differentiation, which further influences inflorescence architecture and increases yield by producing more inflorescence (Jiang et al., 2013). However, the detailed molecular mechanism underlying the dose effect between *SP* and *SFT* remains unknown.

As a post-transcriptional processing event, RNA editing is defined as an RNA product different from its DNA template, which is a supplement to the central dogma (the flow of genetic information from DNA to RNA to protein). RNA editing was first discovered in *Trypanosoma brucei* mitochondria (Benne et al., 1986). In plants, RNA editing (the C-to-U transition) is highly prevalent and restricted within organelles (mitochondria and chloroplasts), which can alter the coding sequences of the organellar transcripts (Rajasekhar and Mulligan, 1993; Gerke et al., 2020; Small et al., 2020). Thus far, despite reports on the genome-wide RNA editing of nuclear transcripts in humans (Chen, 2013) and *Arabidopsis* (Meng et al., 2010), none of these predictions has been verified by gold-standard Sanger sequencing, and few studies have been performed on the involvement of nuclear transcript RNA editing in regulating important traits, such as plant flowering.

In this study, we report the identification of *BOSS* (<u>B</u>alancer <u>o</u>f *<u>S</u>P* and *<u>S</u>FT*) as the causal gene corresponding to our previously identified FIN-related QTL (Ye et al., 2020), *qFIN9* (QTL of FIN on chromosome 9) in tomato (Table S1). *BOSS* has two transcripts (*BOSS-α* and *BOSS-β*) attributed to endogenous RNA editing. *BOSS-β* rather than *BOSS-α* is shown to promote flowering because of their different regulatory patterns with *SP*. This study reveals a dynamic mechanism in floral initiation which confers to the balance of *SP* and *SFT* in tomato.

## RESULTS

### Characterization of Endogenous RNA editing of *BOSS*

An E3 ubiquitin-protein ligase (*Solyc09g010650.1*), which has been reported to regulate flowering in rice (Yang et al., 2015), was preferred as the candidate of *qFIN9. BOSS* contains only one single exon and encodes a 176-amino acid protein with a molecular mass of 16.74 kDa, with an E3 ubiquitin-protein ligase domain present in its C-terminal region (amino acids 78–120) and a transmembrane domain located in its N-terminal region (amino acids 7–26) (Fig. S1A). Sequence analysis showed that the orthologous gene of *BOSS* in *Arabidopsis thaliana* is an *ATL33* gene, which lacks relevant research, with almost all homologous genes present in monocotyledons (Fig. S1B). The expression of *BOSS* is high in seed and opened flower, but low in root and bud (Fig. S1C). Only one polymorphism, Chr09_3978372 (A/T, SNP1) in the 5 ‘UTR of *BOSS*, and no significant difference was detected in the *BOSS* expression between the Low-FIN and High-FIN lines (Fig. S2), suggesting that the two *BOSS* haplotypes are independent of the transcriptional level.

Interestingly, in the individual plants of TS54 (*BOSS*^AA^), we found two transcripts (*BOSS*-α and *BOSS*-β) were distinguished by a SNP (SNP2, C/U at position 457 of RNA sequence, with amino acid being changed from R to W at position 153 of amino acid sequence) (Fig. 1A). As we know, RNA editing has been defined as a single-base difference between DNA and RNA, and the common model of RNA editing in plant organelles is “C>U”, indicating a good match of the phenomenon we found in the nuclear gene *BOSS* for RNA editing. Random primers were used to detect the RNA editing of organelles previously (Wang et al., 2019), but *BOSS* is a nuclear gene decorated with a polyA after transcription. Whether the type of reverse transcription will influence the detection of RNA editing was investigated by using three methods for reverse transcription: only random primers, only oligodT primers, and a combination of the two. The random & oligodT primers were shown to detect the RNA editing of *BOSS* more stably and thus were used in further studies (Fig. S3). To verify the RNA editing in *BOSS*, we sequenced the gDNA and cDNA of *BOSS* in the three individuals of TS54 belonging to *BOSS*^AA^. In Fig.1B, the gDNA sequence of *BOSS* was shown as C at position 457 in TS54, in contrast to the heterozygous peak of C and T for the cDNA sequence at this position.

**Fig. 1.**
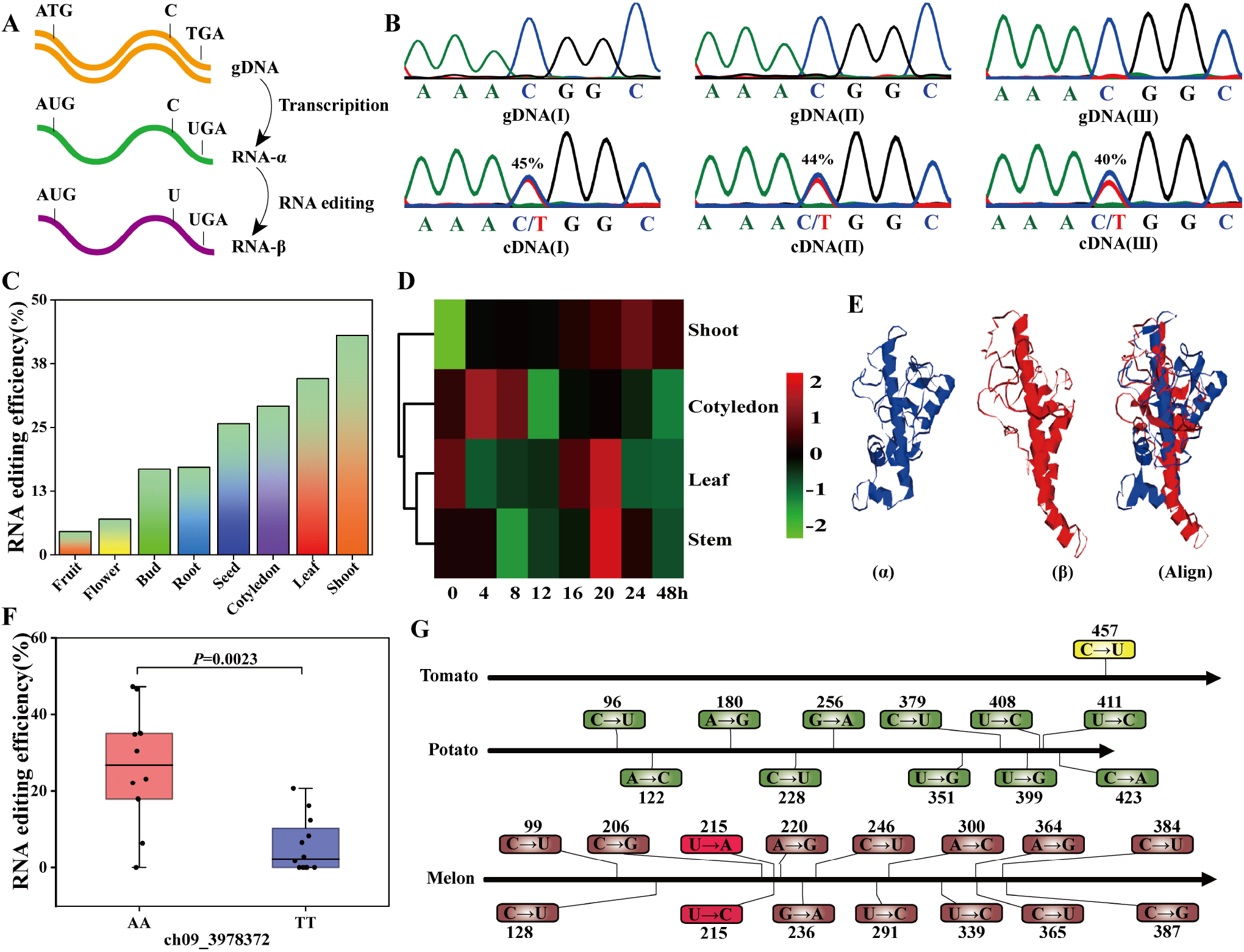
RNA editing of *BOSS*. (**A**) Schematic diagram of *BOSS* RNA editing. *BOSS*-α with C at position 457 is obtained by transcription from gDNA to RNA, and then *BOSS*-β with U at position 457 is formed by RNA-editing. (**B**) The RNA editing efficiency of *BOSS* in TS54. Sequence analysis at RNA-editing site reveals 43% RNA editing efficiency of *BOSS* in TS54, with three independent plants used for the analysis. Color traces: G-black, A-green, T-red, C-blue. (**C**) The RNA-editing efficiency of *BOSS* in different organs of TS54. (**D**) The role of RNA-editing efficiency of *BOSS* in light response. The RNA-editing efficiency of *BOSS* in different organs of TS54 were detected under 48-h continuous light. (**E**) Changes in the three-position structure of BOSS protein due to non-synonymous mutations induced by RNA-editing. (**F**) The RNA-editing efficiency of *BOSS* in the leaves of two *BOSS* haplotypes based on SNP1 (A/T). (**G**) The orthologous genes PGSC0003DMT400023038 and Csa_3g734330 of the *BOSS* gene in potato and melon showed more RNA editing sites than the tomato *BOSS* gene. \**P* < 0.05, \*\**P* <0.01.

The tissue distribution of RNA-editing in *BOSS* was analyzed by using TS54 (*BOSS*^AA^) as an example, which was found to be widely present in the root, stem tip, leaf, cotyledon, flower, bud, seed, and fruit, with the RNA-editing efficiency being highest in the stem tip and lowest in the fruit (Fig.1C). As tomato flowering is strictly regulated by light, we investigated its effect on the RNA-editing efficiency of *BOSS* in cotyledon, stem, leaf and stem tip (Fig.1D). Under light treatment, the RNA-editing efficiency was continuously induced and reached its peak at 24 h in stem shoot, most rapidly induced at 4 h and followed a downward trend in cotyledon, and exhibited a decrease first and then an increase to the peak at 20 h in leaf and stem. In terms of time, RNA-editing efficiency reached its peak most rapidly (at 4 h) in cotyledon (an increase of 8% per hour), 16 and 20 hours earlier than in leaf and stem tip (an increase of 1% per hour), respectively. The results of the light experiments suggest that the earliest receptor organ for the photoinduced RNA-editing efficiency of *BOSS* was cotyledon, followed by leaf, stem and stem tip.

The 3D structure prediction of the BOSS protein showed that RNA-editing leads to the production of *BOSS* genes with different protein structures (Fig. 1E). Moreover, sequence analysis revealed more predicted protein binding sites for BOSS-β than BOSS-α, suggesting the higher activity of the former (Fig. S4A). The similarities and differences between BOSS-α and BOSS-β in the subcellular localization were investigated by creating BOSS-YFP fusion proteins, which were transiently expressed in *Nicotiana benthamiana.* The fluorescent signals of YFP were seen to overlap with those of N-RFP (a marker for the nucleus) and M-RFP (a marker for the plasma membrane), suggesting that both BOSS-α and BOSS-β are located in the nuclear and plasma membranes (Fig. S4B, C).

The RNA editing in *BOSS* and its association with SNP1 were verified by sequencing the gDNA and cDNA of *BOSS* in 23 accessions, with 12 of them representing *BOSS*^TT^ and 11 of them belonging to *BOSS*^AA^. In Fig.1F, *BOSS*^AA^ was seen to have higher RNA-editing efficiency (an average of 43%) than *BOSS*^TT^ (an average of 8.9%), revealing the association of RNA-editing efficiency with SNP1 in 5 ‘UTR.

Whether RNA-editing of *BOSS* is conserved in different lineages was tested by identifying the BOSS orthologous proteins in potato (*Solanum tuberosum*) and melon (*Cucumis melo*), of which more RNA editing types were found than tomato (Fig. 1G). The above results suggest that RNA editing of different complexity may be conserved in different flowering plants.

### *BOSS*-β instead of *BOSS*-α promotes flowering in tomato

The functional differences between *BOSS*-α and *BOSS*-β in tomato flowering were evaluated by generating two overexpression constructs containing allele *BOSS*-α and *BOSS*-β and introducing them into TS221 (*BOSS*^TT^ with low RNA-editing) and ZY3 (*BOSS*^AA^ with high RNA-editing), respectively. The *BOSS* RNA levels showed a significant increase in the three independent *BOSS*-β overexpression lines in T_1_ generation (Fig. 2A), with the *BOSS*-β transcript being 100% in the *BOSS*-β overexpression lines (Fig. 2B). Compared with TS221, the three *BOSS*-β overexpression lines had lower FIN and FSIN (<u>f</u>irst to <u>s</u>econd <u>i</u>nflorescence <u>n</u>ode) (Fig. 2 C-E). The early flowering of *BOSS*-β overexpression lines was further evaluated by microscopic observation of paraffin sections and SAM scanning electron micrographs from wild-type tomato and *BOSS*-β overexpression lines (Fig. 2F, G). The FM (flower meristem) of the primary meristem was observed to occur earlier in the *BOSS*-β overexpression lines than in the wild-type tomato, leading to early flowering. Moreover, the lower FIN and FSIN increased the flower trusses, thereby increasing the yield per plant in the *BOSS*-β overexpression lines (Fig. 2H, I). Interestingly, the overexpression of *BOSS*-β led to the self-pruning of a few transgenic plants (Fig. S5), suggesting that *BOSS* may regulate self-pruning by some dosage interaction with other related genes. Conversely, the overexpression of *BOSS*-α failed to affect the differentiation of FIN, FSIN or FM in ZY3 (Fig. S6). The results of the transgenic experiments strongly suggest that *BOSS*-β rather than *BOSS*-α functions as a positive regulator of tomato flowering.

**Fig. 2.**
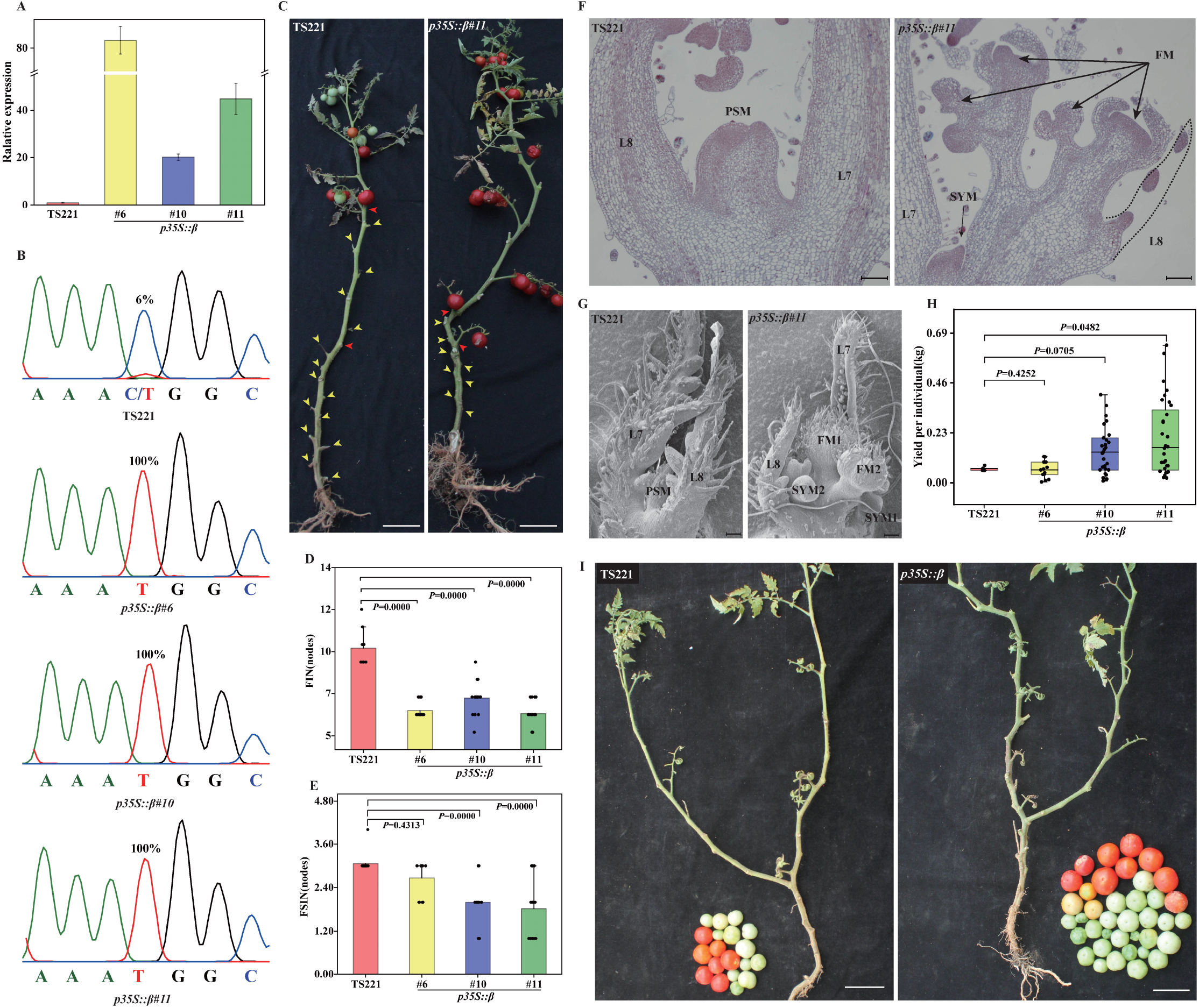
Elevated *BOSS*-β levels contribute to early flowering in tomato. (**A**) The relative expression of *BOSS* in *BOSS*-β overexpression transgenic tomato lines. (**B**) Sequence analysis at RNA-editing site reveals 100% RNA editing efficiency of *BOSS*-β overexpression versus 6% in TS221, with three independent plants used for the analysis. Color traces: G-black, A-green, T-red, C-blue. (**C**) Primary inflorescences (red arrowheads) and side branches (yellow arrowheads) from TS221 (left) and *BOSS*-β overexpression line (right) with early flowering. (**D**, **E**) Quantification and comparison of FIN (first inflorescence node) (**D**) and FSIN (node between first and second inflorescence) (**E**) in TS221 and *BOSS*-β overexpression lines. (**F, G**) Stereomicroscopic (**F**) and scanning electron microscopic (**G**) analysis of the primary shoot apical meristems from TS221 and *BOSS*-β overexpression lines. (**H**, **I**) Quantification (**H**) and images (**I**) of yields from TS221 and *BOSS*-β overexpression lines. Scale bars: 5 cm (C), 100 μm (F), 100 μm (G) and 5 cm (I). All data in the graphs represent means ± SD (*n* = 3); \**P* < 0.05, \*\**P* <0.01.

### *BOSS*-β promoting flowering by inhibiting the expression of SP

How different *BOSS* transcripts regulate tomato flowering was explored by RNA-seq analysis of ZY3/*BOSS*-α-OE and TS221/*BOSS*-β-OE lines. Differential gene expression analysis identified the up- and down-regulation of 36 and 33 genes, respectively, in TS221/*BOSS*-β-OE lines (fold change >2; FDR < 0.05), in contrast to the up- and down-regulation of only 7 and 2 genes, respectively, in ZY3/*BOSS*-α-OE line (Table S2, Fig. S7A, B). Additionally, GO enrichment analysis of the upregulated genes in TS221/*BOSS*-β-OE reported ‘MADS-box transcription factor’ as the top term and included the known flowering-related gene FLC-like (Solyc12g087810) (Fig. S7C). The upregulation of these three MADS-box transcription factors (*Solyc12g087810, Solyc12g087820, Solyc12g087830*) was verified by quantitative RT-PCR (RT-qPCR) (Fig. S7D-F).

Moreover, to analyze the dynamic expression changes of flowering related genes in *BOSS* over-expression lines, we performed quantitative RT-PCR (qRT-PCR) at the different stages of seedling development (Fig. 3). In agreement with the early flowering phenotype, we found that *BOSS*-β inhibited the expression of anti-florigen *SP* (−7.5 folds) at 5^th^ leaf stage to promote flowering, whereas the expression of *SP* (−8.6 folds) was promoted at 6^th^ leaf stage in *BOSS*-α over-expression line (Fig. 3A). The expression of florigen *SFT* was not significantly affected in both *BOSS*-α and -β over-expression lines (Fig. 3B). These results indicated that *BOSS*-β regulated flowering of tomato by regulating *SP* expression.

**Fig. 3.**
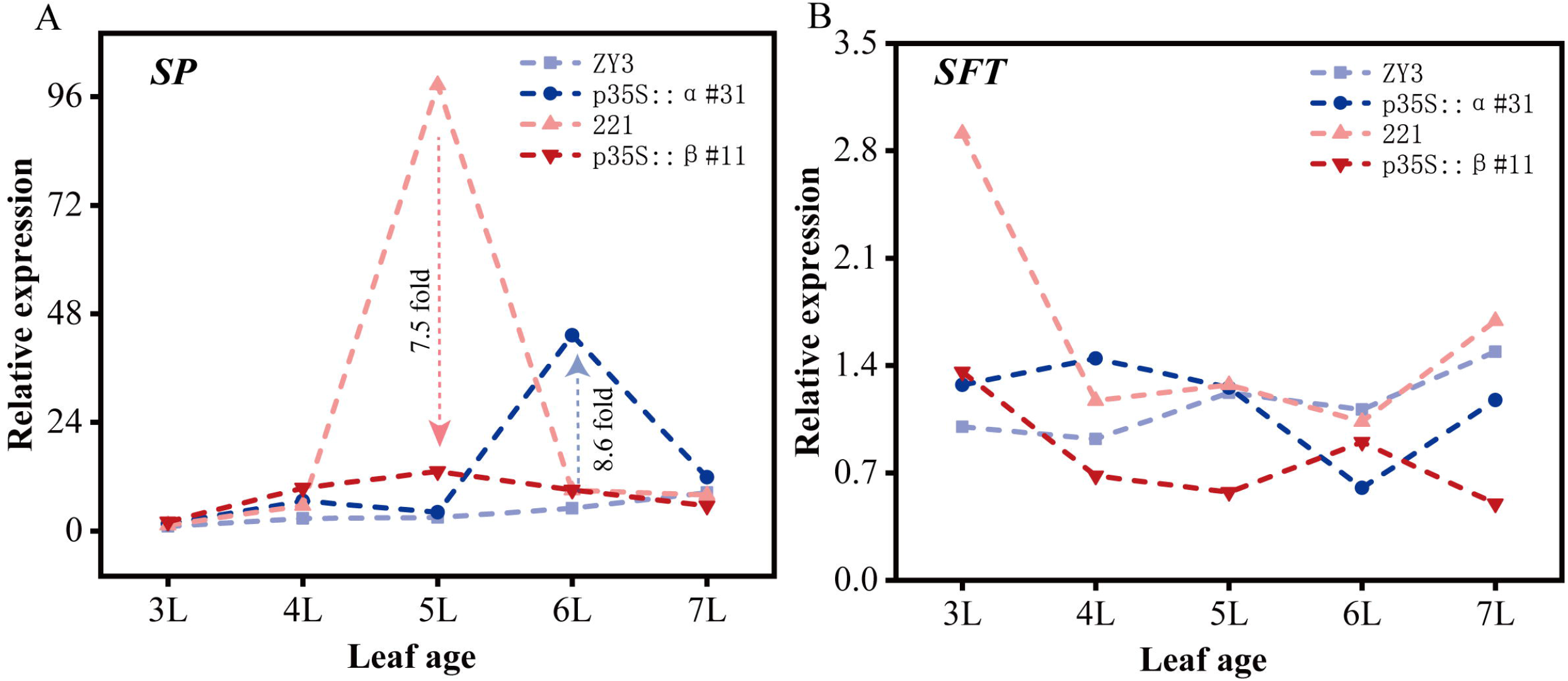
Relative expression dynamics for *SP* and *SFT* in stem apical at the different stages of seedling development. (A) Relative expression dynamics of *SP* in stem apical at the different stages of seedling development. 3L-7L means from 3^th^ leaf state to 7^th^ leaf state. At 5^th^ leaf stage, compared with WT, the expression of *SP* decreased 7.5 times in *BOSS*-β overexpression lines, while overexpression of *BOSS*-α increased the expression of *SP* gene by 8.6 times at 6^th^ leaf stage. (**B**) Relative expression dynamics of *SFT* in stem apical at the different stages of seedling development.

## DISCUSSION

The genome editing events of plant organelles are mainly investigated by bioinformatics and experimental approaches (Tang and Luo, 2018; Yan et al., 2018; Small et al., 2020), while the RNA editing of nuclear transcripts is only studied using bioinformatics predictions in plant (Meng et al., 2010). In this study, we identified the RNA editing of a nuclear gene *BOSS* by Sanger sequencing method which was considered as the gold standard for identification of RNA editing (Zhao et al., 2019). Interestingly, except for TS-225(C/T heterozygosity), all the re-sequenced accessions showed C/C homozygou of the RNA-editing site at DNA level (https://solgenomics.net/organism/Solanum_lycopersicum/tomato_360), suggesting that the variation of this site in natural population may trend to transit from DNA level to RNA level.

Except for tomato, the RNA-editing also widely identified in other species, such as potato and melon (Fig. 1G). Like RNA-editing of *BOSS* regulate *SP* in tomato, the more extensive editing types of *BOSS* orthologous in potato may be of great significance for tuber development which was regulated by *StSP6A*, the orthologous of tomato *SP* (Abelenda et al., 2014). And the significance seems to be greater due to a dual editing site of *BOSS* orthologous gene in melon (Fig. 1G).

In summary, our results confirm the existence of RNA editing of the nuclear transcript *BOSS* (Fig 1). Additionally, we constructed a sophisticated regulatory pathway that the natural variation of SNP1 located in the palindromic sequence of *BOSS* 5 ‘UTR leads to different RNA-editing efficiency, which in turn affects the transcripts of *BOSS*-α and -β. Finally, different transcripts of *BOSS* regulate tomato flowering by affecting the expression level of *SP* at seedling stage, *BOSS*-β decreased the expression of *SP* but *BOSS*-α increased the expression of *SP* (Fig. 4). Although the expression of *SP* both altered in *BOSS*-α and -β overexpression lines, the change expression phase, vital to stem apical meristem fate determinate, was different (5^th^ leaf stage for *BOSS*-β and 6^th^ leaf stage for *BOSS*-α) (MacAlister et al., 2012). The new knowledge about RNA-editing of tomato flowering will facilitate genome-wide research on the regulatory role of RNA editing in various biological processes.

**Fig. 4.**
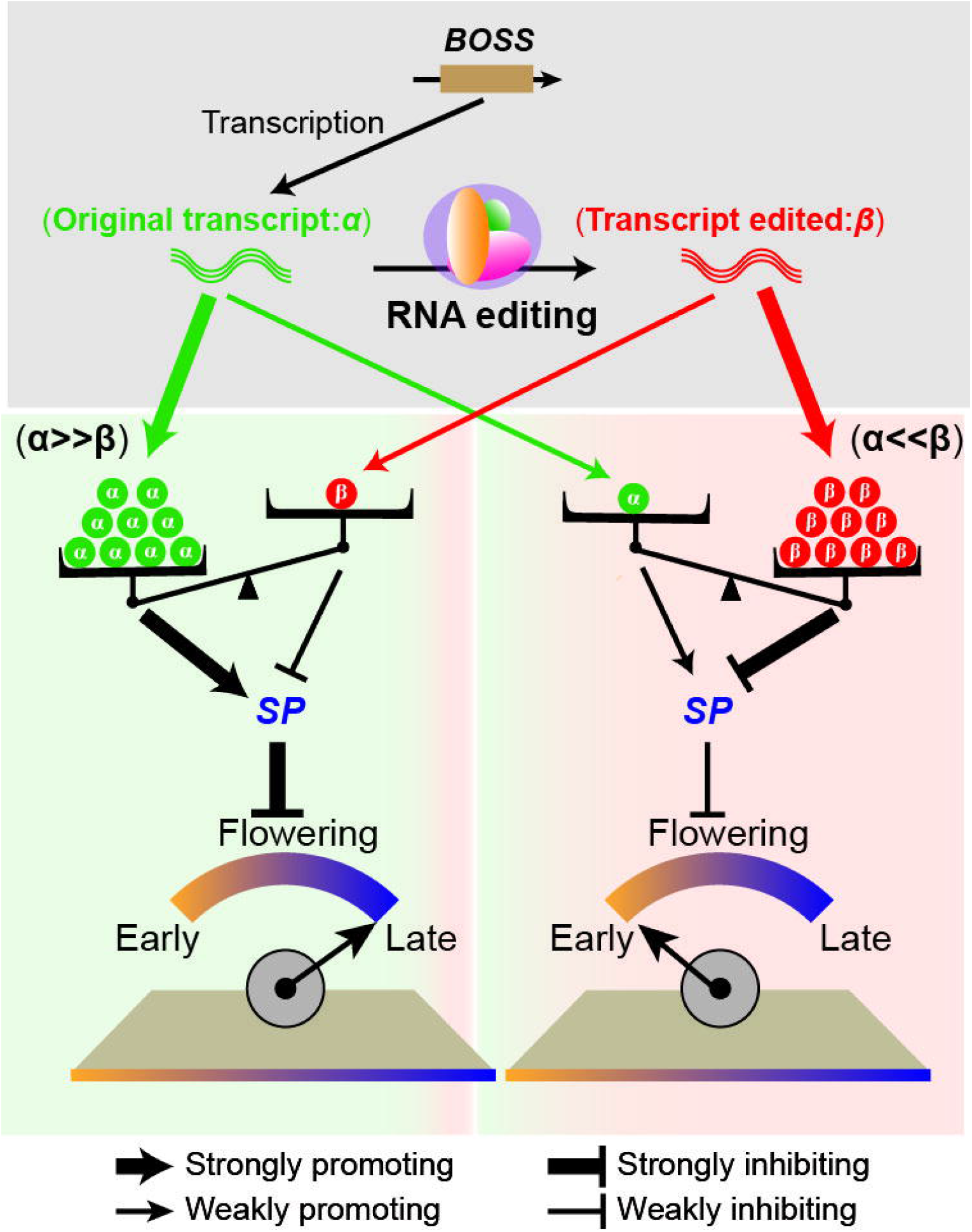
Proposed model of *BOSS* function as a determinant of FIN and regulating tomato flowering. Different transcripts of *BOSS* regulates tomato flowering by affecting the expression level of *SP* at seedling stage. When the transcripts of *BOSS*-α were more than *BOSS*-β, it can promote the expression of *SP* and inhibit flowering. On the contrary, the expression of *SP* was reduced when the transcripts of *BOSS*-α were more than *BOSS*-β, leading to the later flowering.

## METHODS

### Phylogenetic analysis of *BOSS* gene family

The amino acid sequences of *BOSS* and its homologues were aligned using the CLUSTALX (version 2.1) software. The neighbor-joining tree was constructed using aligned full-length amino acid sequences (MEGA6), with bootstrap values from 1,000 replicates indicated at each node. Bar = 0.1 amino acid substitutions per site. The tree was visualized by the online tool Interactive Tree of Life (iTOL).

### Light treatment

TS54 plants (high RNA-editing efficiency) were grown in plastic pots in the greenhouse under a 16-h light and 8-h dark at 25°C. At the seedling age of three leaves and one hear, the plants were treated with continuous light and samples of different tissues were collected at 0 h, 4 h, 8 h, 12 h, 16 h, 20 h, 24 h and 48 h post light treatment to determine the RNA-editing efficiency of *BOSS* in different tomato tissues.

### RNA isolation and gene expression analysis

To analyze the dynamic expression changes of *SP/SFT* at different developmental stages, tomato seedlings were planted and cultured as above, and shoot tip tissues were collected from 3th leaf stage to 7th leaf stage, respectively. Total RNA was isolated from tomato using TRIZOL reagent (Invitrogen, USA) according to the manufacturer’s instructions. The cDNAs were synthesized from 1μg total RNA using HiScript^®^II Reverse Transcriptase (Vazyme, China) following the manufacturer’s protocol. Gene expression was investigated by quantitative real-time PCR (qRT-PCR) as previously described (Ye et al., 2019). The actin gene (*Solyc11g008430*) was used as an internal standard and the relative expression of specific genes was measured using the cycle threshold (Ct) 2^−ΔΔCt^ method.

### RNA-Seq analysis

At the seedling age of six weeks, the shoot tip tissues from TS221, *BOSS*-α-OE and *BOSS*-β-OE were frozen in liquid nitrogen and kept at −80°C until further use. An RNA sample was obtained from three plants of each genotype, with three biological replicates (3 × 3 = 9 plants) for each genotype. Total RNAs were extracted using the same method as described above and then sent to the Berry Genomics Company (Beijing, China), where the libraries were constructed and sequenced using single-ended sequencing of Illumina Novaseq6000. The sequencing data can be accessed at the Sequence Read Archive (SRA ID: PRJNA673993). The reference genome sequences (SL2.50 version) of *S. lycopersicum* were downloaded from the SOL Genomics Network database (http://solgenomics.net/organism/Solanum_lycopersicum/genome). Here, genes with a *P* value < 0.01 and a log2 ratio > 1.0 or < 0.5 were considered differentially expressed. Sequence analysis and differential expression analysis were performed with a method provided by Novogene. More detailed information is provided on the Novogene website (www.novogene.com).

### Analysis of RNA editing efficiency in *BOSS*

The coding sequence of *BOSS* containing RNA editing site was amplified by PCR with cDNA as the templet and the specific primers (Table S3) using Phanta Super-Fidelity DNA Polymerase (Vazyme, China). The PCR was conducted as follows: 95 °C for 3 min, 34 cycles of 95 °C for 15 s, 56 °C for 15 s, 72 °C for 30 s, and 72 °C for 5 min. The PCR products were used as templates for Sanger DNA sequencing (Sequencing was performed by Tianyi Huiyuan Bioscience and Technology, Wuhan, China). The “C” to “T” ( C to U in RNA) editing efficiency was measured by the relative height of the peak of the nucleotide in sequence chromatograms and calculated by the height of “T” / (height of “T” and “C”) as previously described (Zhao et al., 2019).

### Subcellular analysis

Using the cDNA of TS221 (low RNA-editing efficiency) and TS54 (high RNA-editing efficiency) as templates, the coding sequences of *BOSS*-α and *BOSS*-β without the stop codon were amplified by PCR using the primer sequences shown in Table S3, then cloned into the expression vector p101YFP under the control of the CaMV35S promoter by homologous recombination (ClonExpress II One Step Cloning Kit, Vazyme, China). CaMV35S:BOSS-α/β-YFP vector as well as cell nucleus (nucleolus) marker CaMV35S:N-RFP (AtNM, AT4G25630.1) (Degenhardt and Bonham-Smith, 2008), plasmid membrane/nuclear membrane marker CaMV35S:*CBLn1*-RFP (Batistic et al., 2010) and the control YFP vector (positive control) were transformed into *Agrobacterium tumefaciens* strain GV3101 and co-infiltrated into leaves of *N. benthamiana* with the suspension as previously described. After 48-h incubation at 25 °C, the tobacco leaves were used for YFP and RFP observation using Leica Confocal software.

### Gene cloning and vector construction

For the overexpression construct, the 1032bp fragment containing the 5’UTR (including 8T8A) and CDS region of *BOSS* was amplified from the genomic DNA of TS221 (9T7A_5’UTR_-α_CDS_ is the main transcript type) and TS-226 (8T8A_5’UTR_-β_CDS_ is the main transcript type) to obtain 9T7A_5’UTR_-*BOSS*-α_CDS_ and 8T8A_5’UTR_-*BOSS*-β_CDS_ fragments by point mutation. The PCR products were inserted into pDONR221 using the BP enzyme (Invitrogen, USA), and then incorporated into the destination vector pMV3 (driven by the cauliflower mosaic virus (CaMV) 35S promoter) using the LR recombination reaction (Invitrogen, USA). All the recombinant constructs were transformed into the *Agrobacterium tumefaciens* strain C58 by electroporation and subsequently transformed into the tomato genome *via* explants of cotyledon. To eliminate the influence of *SP* locus in chromosome 6, the background materials TS221 (*SP/SP*, 9T7A with lower RNA editing efficiency) and ZY3 (*SP/SP*, 8T8A with higher RNA editing efficiency) were selected to transform 8T8A_5’UTR_-*BOSS*-β_CDS_ and 9T7A _5’UTR_-*BOSS*-α_CDS_, respectively.

### Data availability

Data supporting the findings of this work are available within the paper and its Supplementary Information.

## Acknowledgments

This study was supported by the National Natural Science Foundation of China (31991182 and 31872118), the National Key Research and Development Program of China (2017YFD0101902) and CARS-23-A-03.

## Author Contributions

Z. Y., H. L., H. H., J. Z., B. O., W. W., C. Y., X. W., C. Y. and J. Y. designed research and managed the project; W. W., C. Y., Q. X., J. S. and L. C. performed research; W.W., J. Y., H. Y., C. L., Y. L., T.W. and Y.Z. analyzed data; J.Y., W.W. and Z.Y. wrote and modified the manuscript.

## SUPPORTING INFORMATION

**Fig. S1. Characterization of *BOSS*.** (**A**) *BOSS* gene diagram, including 5’UTR, CDS and 3’UTR. (**B**) Phylogenetic tree of *BOSS* and its homologous genes in different species. (**C**) The transcript levels of *BOSS* in different tomato organs: Ro, root; FB, Flower bud; Le, leaf; Fr, Fruit; St, stem; Fl, flower; Se, Seed.

**Fig. S2. The relative expression of *BOSS* in the stem apices of 23 selected accessions (12 SNP1^AA^ accessions and 11 SNP1^TT^ accessions).** Significant difference was calculated by *t*-test: \*\**P*<0.01.

**Fig. S3. The influence of different reverse transcription methods on the detection accuracy of RNA editing efficiency.** Three reverse transcription methods were used: only random primers, only oligodT primers, and combination of the two.

**Fig. S4. Effect of BOSS RNA-editing on protein interaction sites (A) and subcellular localization (B, C)**. (**A**) Prediction of changes in the protein RNA interaction sites by BOSS RNA-editing (http://pridb.gdcb.iastate.edu/RPISeq/). (**B, C**) Subcellular co-localization of transiently expressed *BOSS*-α/β-YFP fusion protein with nuclear membrane marker (**B**) and plasmid membrane marker (**C**) in *N. benthamiana* leaves. Bar =10 μm.

**Fig. S5. Overexpression of *BOSS*-β leads to determinate sympodial apices.** The percentages of determinate plants in the three transgenic lines are shown in Spring 2016 (**A**) and Spring 2017 (**B**).

**Fig. S6. Increasing the expression of *BOSS*-α does not affect tomato flowering.** (**A**) The relative expression of *BOSS* in *BOSS*-α overexpression transgenic tomato lines. (**B**) Sequence analysis at RNA-editing site reveals 0% RNA editing efficiency of *BOSS*-α overexpression versus 72% in ZY3, with three independent plants used for the analysis. Color traces: G-black, A-green, T-red, C-blue. (**C**) Primary inflorescences (red arrowheads) and side branches (yellow arrowheads) from ZY3 (left) and *BOSS*-α overexpression line (right) with no flowering change. (**D, E**) Quantification and comparison of FIN (first inflorescence node) (**D**) and FSIN (node between first and second inflorescence) (**E**) in ZY3 and *BOSS*-α overexpression lines. Scale bar=5 cm (C). All data in the graphs represent means ± SD (*n* = 3).; \**P* < 0.05, \*\**P* < 0.01.

**Fig. S7. RNA-seq analysis of differentially expressed genes in the shoot apical meristem of** *BOSS*-α/β **overexpression lines versus wild type.** (**A, B**) Volcano maps for the differentially expressed genes in *BOSS*-β overexpression line (**A**) and *BOSS*-α (**B**) overexpression line versus the wild type. (**C**) GO enrichment analysis of differentially expressed genes in *BOSS*-β overexpression line versus wild type. (**D-F**) The relative expression of three flowering-related MADS-box transcription factors was up-regulated in *BOSS*-β overexpression transgenic tomato lines.

**Table S1. Genes within 50 kb of the SNP with high relation to FIN.**

**Table S2. Significantly differentially expressed genes (log2(FC)>1 or <-1) in SAM of *BOSS* overexpression line (p35S::β and p35S::α) versus wild type.**

**Table S3. Primers used in this study.**

