## Supplementary figures and images for "Endogenous RNA editing of a nuclear gene *BOSS* triggers flowering in tomato"

### Supplementary Figures

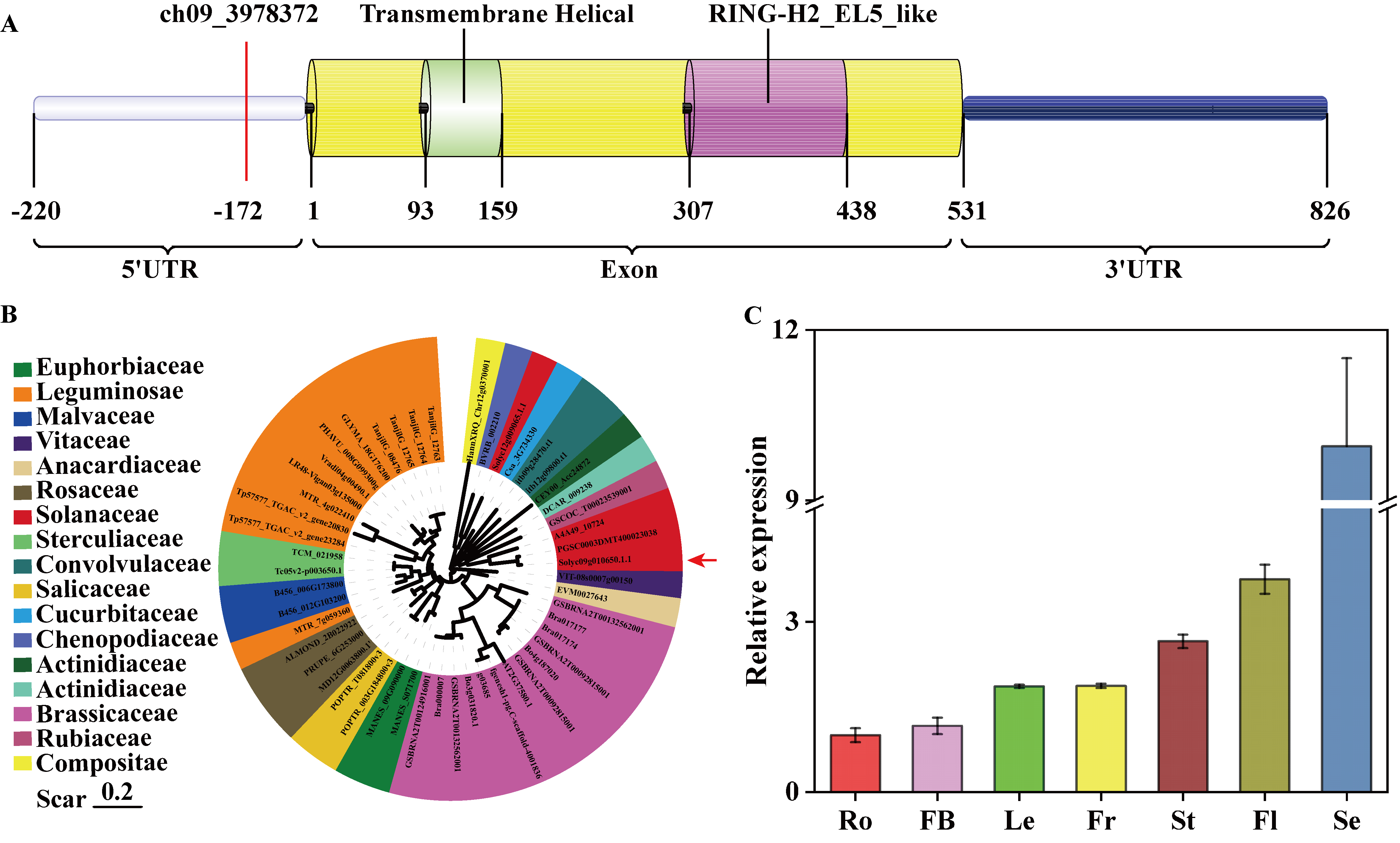

### Supplementary Figures

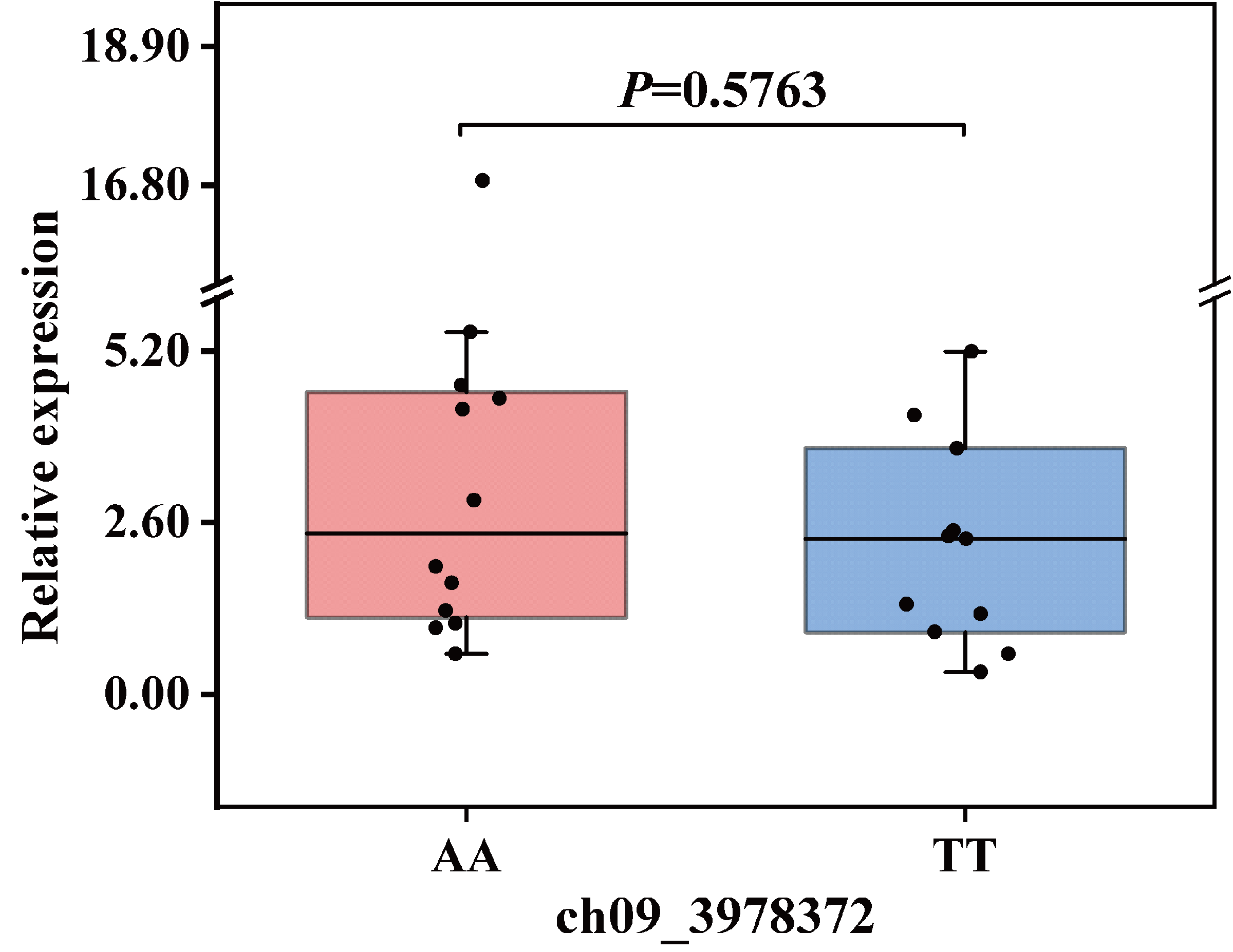

### Supplementary Figures

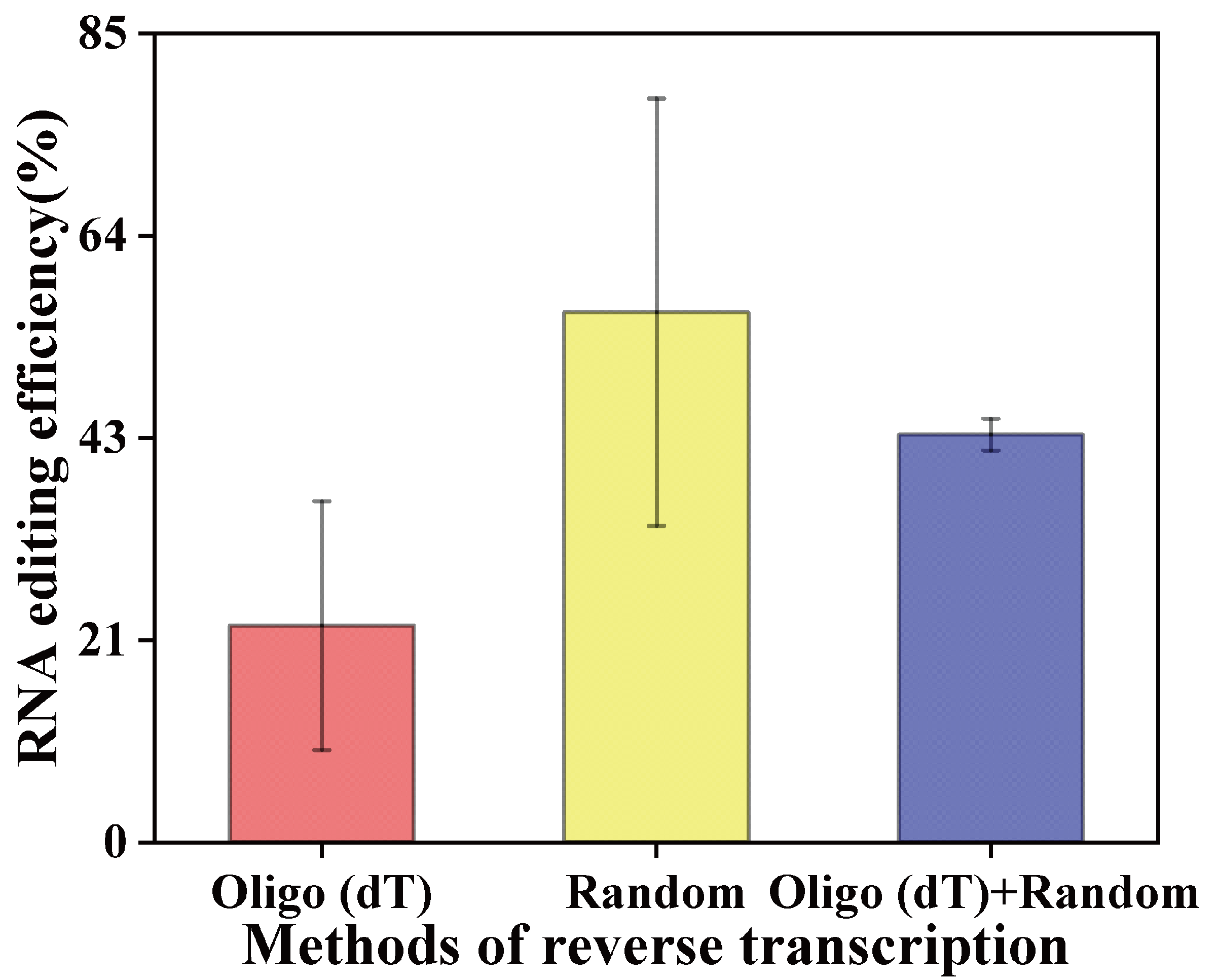

### Supplementary Figures

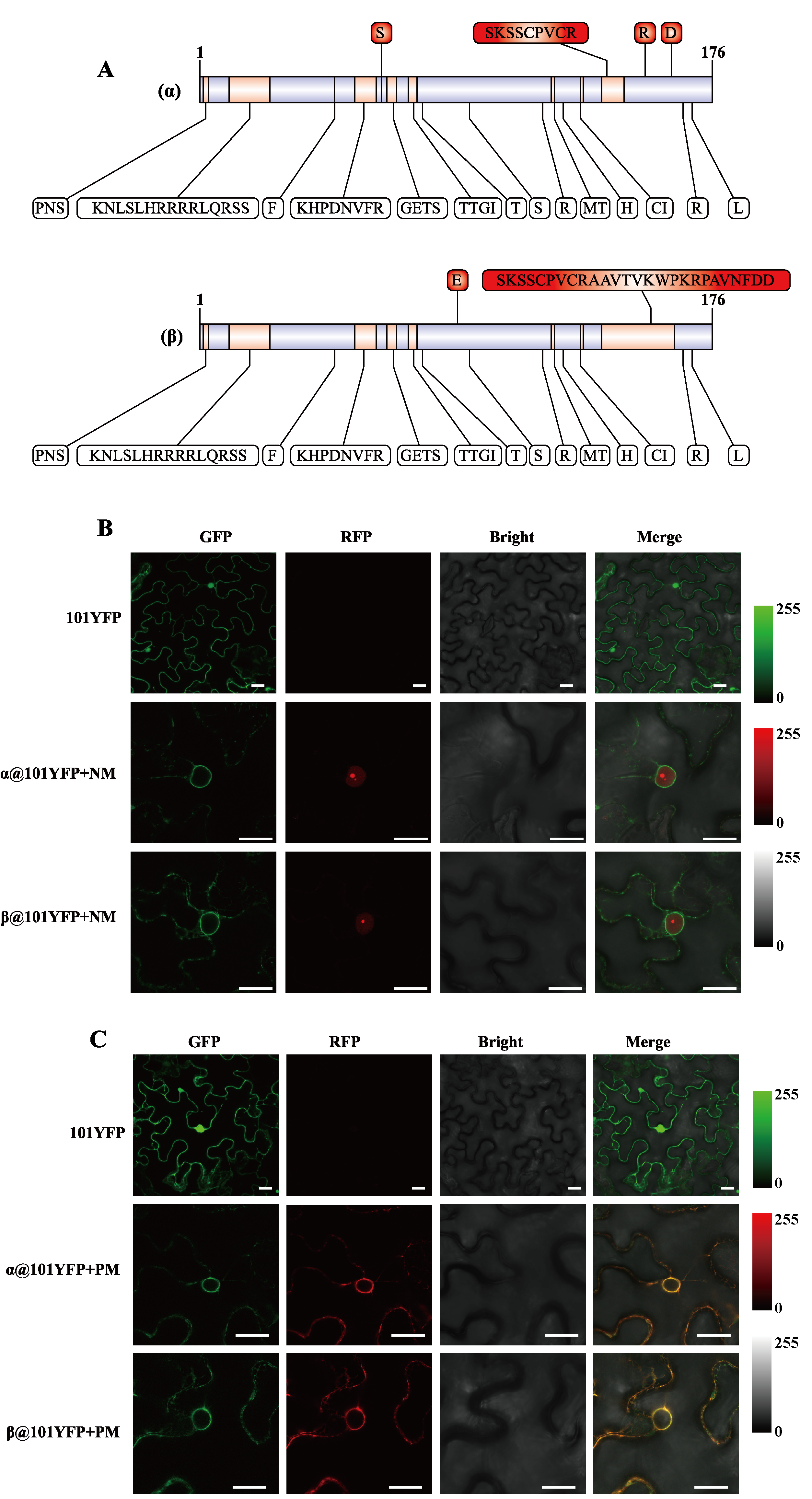

### Supplementary Figures

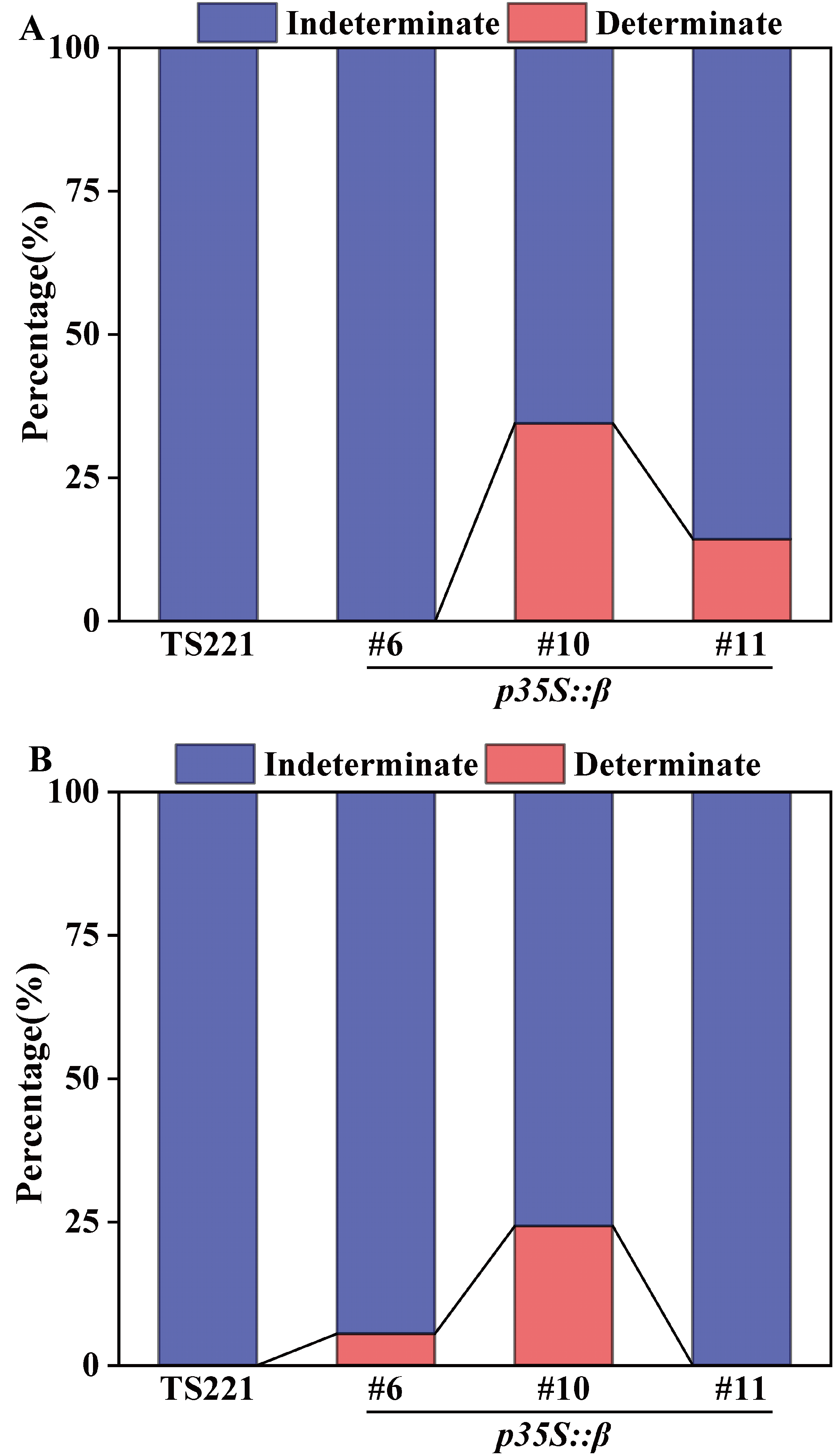

### Supplementary Figures

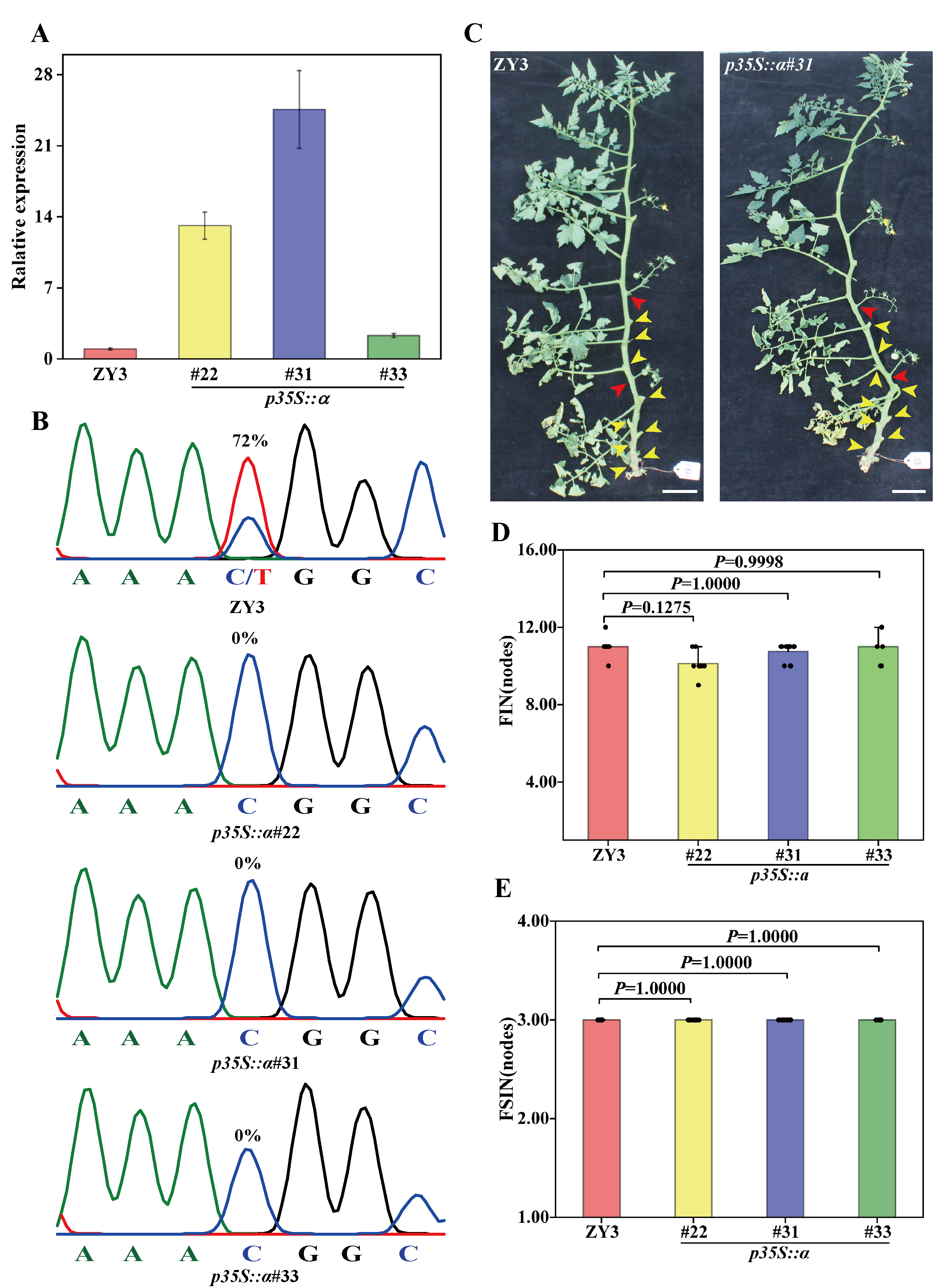

### Supplementary Figures

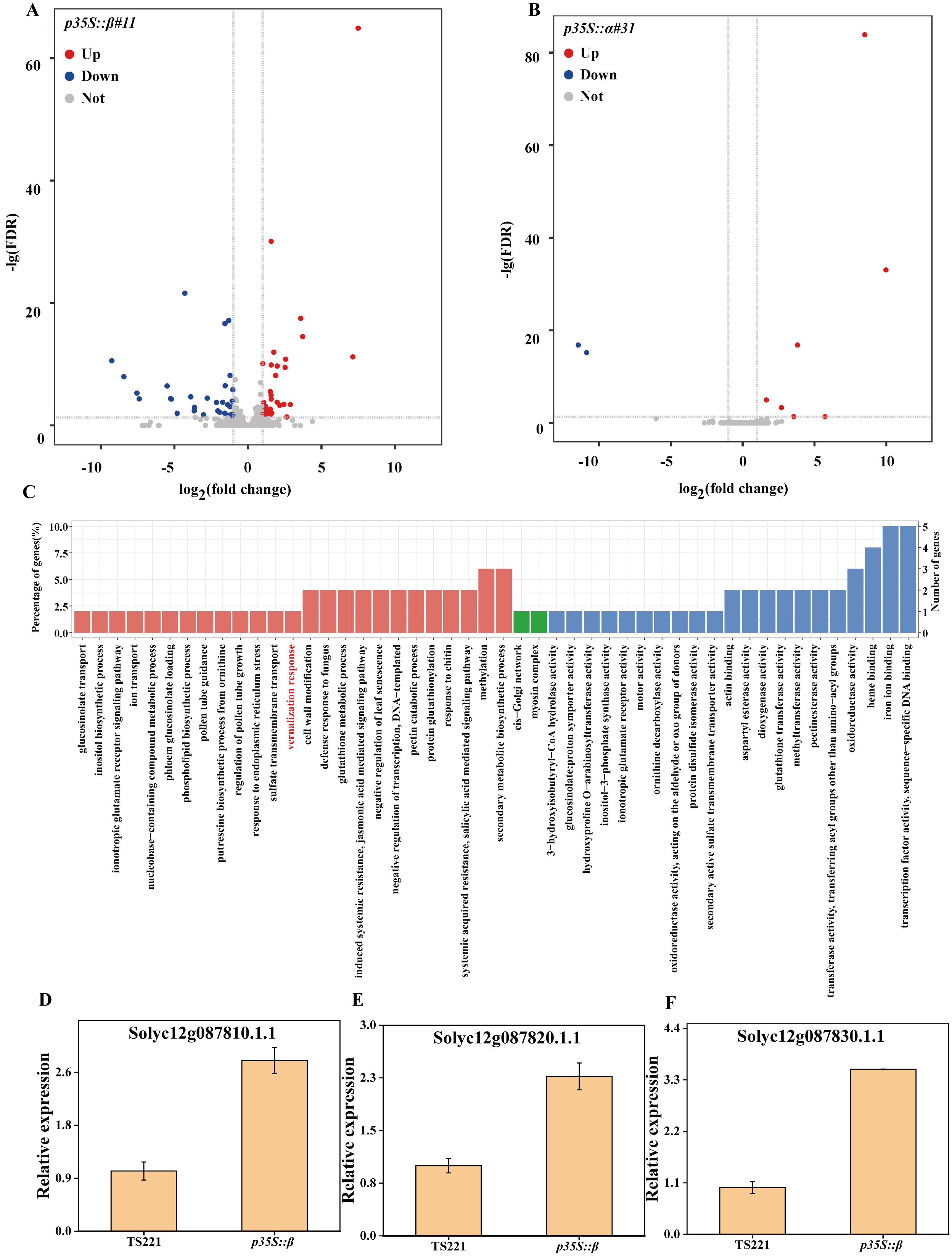
